# Effect of methanol fixation on single-cell RNA sequencing data

**DOI:** 10.1101/2020.05.21.109975

**Authors:** Xinlei Wang, Lei Yu, Angela Wu

**Affiliations:** Division of Life Science, Hong Kong University of Science and Technology, Clear Water Bay, NT, Hong Kong, HKSAR China

**Keywords:** Single Cell RNA-seq, Methanol fixation, Smarts-seq2, Drop-seq

## Abstract

Single-cell RNA sequencing (scRNA-seq) has led to remarkable progress in our understanding of tissue heterogeneity in health and disease. Recently, the need for scRNA-seq sample fixation has emerged in many scenarios, such as when samples need long-term transportation, or when experiments need to be temporally synchronized. Methanol fixation is a simple and gentle method that has been routinely applied in scRNA-seq. Yet, concerns remain that fixation may result in biases which may change the RNA-seq outcome. We adapted an existing methanol fixation protocol and performed scRNA-seq on both live and methanol fixed cells. Analyses of the results show methanol fixation can faithfully preserve biological related signals, while the discrepancy caused by fixation is subtle and relevant to library construction methods. By grouping transcripts based on their lengths and GC content, we find that transcripts with different features are affected by fixation to different degrees in full-length sequencing data, while the effect is alleviated in Drop-seq result. Our deep analysis reveals the effects of methanol fixation on sample RNA integrity and elucidates the potential consequences of using fixation in various scRNA-seq experiment designs.

## Introduction

Since its emergence, single-cell RNA-seq (scRNA-seq) has revolutionized many biological fields due to its high resolution in deciphering tissue heterogeneity (1). The mRNA input from one cell is quite little, thus it leads to more dropout in gene detection compared with bulk RNA-seq (2). During single-cell library preparation, the reverse-transcription (RT) step is crucial since any RNA molecules not captured in this step will forever be lost, and any biases in this step will be amplified downstream, severely affecting the inference of biological signal. For these reasons, it is of utmost importance to preserve the biological sample as much as possible to yield a high-quality transcriptome and a successful scRNA-seq experiment.

For projects including long-distance transportation of samples, cells or tissues may suffer the loss of viability from physical impact during transport or improper storage conditions. In some cases, sample preservation methods are required to allow more flexible experimental designs; specifically, it can help to store samples collected from different experimental conditions or time points and enable them to be consolidated (3). Besides, researchers may also be interested in specific biological states that in some tissues may become altered as specific pathways can be activated by in vitro processing (4).

Fixation has been widely utilized for the preservation of biological samples from postmortem decay. Various fixation protocols that use different chemicals have been developed for different purposes and applications, each method having their pros and cons, partially due to their different fixation mechanisms (5-7). To preserve the desired biological features of tissues or cells, different fixatives play different roles depending on the desired features to be preserved. Crosslinking fixatives, such as formaldehyde, work by creating covalent chemical bonds between proteins in tissues, thereby stopping all enzymatic and macromolecular function in the tissue. This causes a complete arrest of all cellular activity, including cell apoptosis and molecular degradation; most macromolecules are even locked in the spatial position they were in at the time of fixation so that spatial relationships within the cell are also preserved. Formaldehyde specifically fixes tissues by cross-linking primarily the residues of the basic amino acid lysine in proteins, and is an ideal fixative for immunohistochemistry (IHC) (8); as all macromolecules are cross-linked, this kind of fixation offers the benefit of long-term storage and allows good tissue penetration by dyes and other small molecule chemicals required for downstream processing in IHC (9). Another cross-linking fixative, PFA, can anchor soluble proteins to the cytoskeleton and lends additional rigidity to the tissue (10). The FRISCR protocol based on PFA fixation can even integrate fluorescent dye staining, which allows researchers to apply FACS analysis on this type of fixed sample and sort specific cellular subpopulations for further sequencing analysis (11). This protocol is not, however, suitable for adaptation to high throughput scRNA-seq as it requires a reverse crosslinking step that can only be performed in tubes and is not compatible with most microfluidic scRNA-seq library preparation workflows.

Alcohol fixatives, such as ethanol and methanol, work by dehydration, causing proteins to denature and precipitate in-situ (12). As such, the cellular structure will be damaged since the dehydrated environment changes protein conformation. Therefore, alcohol fixation alone is not ideal for preserving samples for imaging, but it is useful for nucleic acid preservation. Compared with fixation approaches used in histology, nucleic acid preserving methods for sequencing do not require the integrity of structural proteins, instead, they aim to prevent DNA or RNA from degradation. Methanol fixation has been widely utilized for its ease of operation and robust performance in preserving nucleic acids (13-14). The dehydration effect can be reversed with a single, simple rehydration step, which can easily be incorporated into scRNA-seq workflows at the sample preparation step, with subsequent processing steps for cDNA library construction carried out normally without any additional changes (15). Although methanol can be largely removed by PBS buffer washing to avoid contamination of downstream reactions, substantial changes occur in cells upon fixation due to dehydration. The cellular structure becomes damaged and normal cell functions are compromised due to loss of normal lipid and protein structure; how these changes affect the transcriptome and whether they will influence the sequencing profile remains understudied. In this study, we comprehensively evaluated the effect of methanol fixation on single-cell RNA-seq results. We performed the analysis at gene and transcript levels and observed both similarities and inconsistencies between the transcriptomic profiles of live and fixed cells. Although it is often assumed that fixation-associated RNA degradation is the main reason for the discrepancies between live and fixed transcriptomic profiles, our results indicate the incomplete reverse transcription of mRNAs with more complex secondary structures during the library preparation step may be another important cause of the observed discrepancies.

## Result

### Methanol fixation does not affect nucleic acid integrity and preserves cell-to-cell similarities consistent with scRNA-seq technical variability

First, we wanted to determine whether there is any obvious degradation of RNA or changes to the transcriptomic profile caused by methanol fixation. To do so, we performed methanol fixation on two cell lines, HCT-116 and HepG2, such that any cell-type specific fixation effects can also be observed and compared across cell types (Figure 1A; for within cell-type comparisons, the result of the HCT-116 cell line is shown here for illustrative purposes. Results are consistent for both cell lines studied (Supplementary Figure S1-S5)). For both cell lines, we prepared RNA-seq libraries from live cells, as well as from fixed cells that were stored in methanol for one-week. We measured the size of single-cell cDNA libraries (Figure 1B) and noted that although no significant change in fragment size distribution was observed for fixed cells, there is a slight decrease in the quantity of cDNA in the 1500bp-2000bp. This result shows that fixation can largely preserve the RNA integrity such that high-quality cDNA can be obtained without severe degradation.

**Figure 1.**
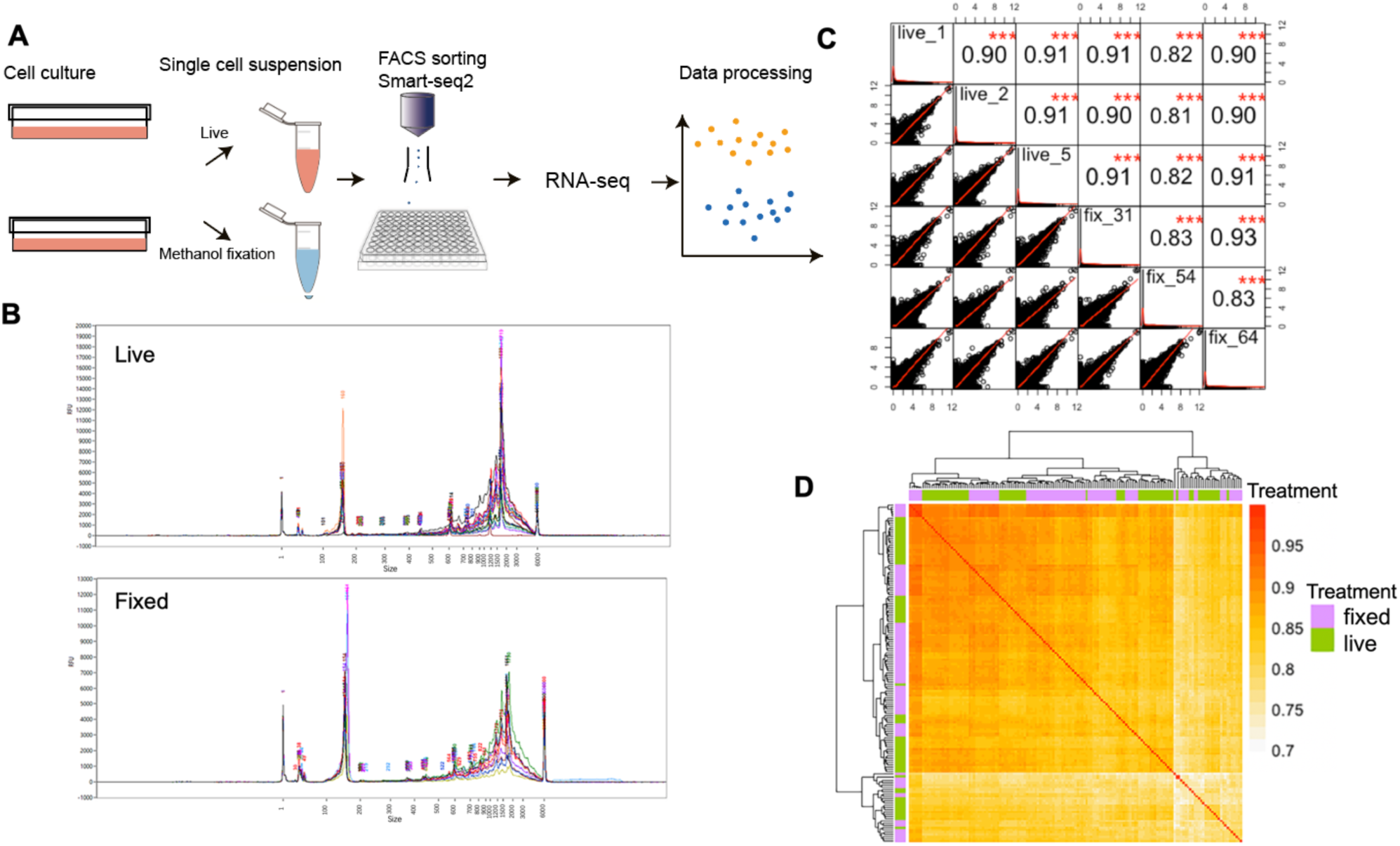
Basic evaluation of fixation effect on sequencing data. **(A)** Workflow and experimental scheme. **(B)** Size distributions of cDNA libraries. Traces from single-cell libraries were merged to obtain a general pattern for live (top) and fixed (bottom) samples. Although the intensity of the ∼1500bp peak is diminished in fixed cells, there is no visible degradation. **(C)** Correlation matrix showing the transcriptome similarity of cells randomly chosen from live and fixed samples. The upper triangle of the matrix shows the Pearson correlation coefficient and the bottom triangle visualized correlation trend. Correlations are consistently high for both inter- and intra-treatment comparisons of live vs. fixed. There is no obvious bias revealed by measuring correlation between single-cell transcriptomes for all pairwise comparisons. **(D)** Correlation factors of all single cells were calculated pairwise and clustered by Euclidean distance. Correlations are consistently high for both inter- and intra-treatment comparisons of live vs. fixed (R^2^ >0.7). The mixed annotation bar indicates the transcriptome similarities do not distinguish cell treatments during sample preparation.

Next, we performed a more detailed bioinformatic analysis to compare the transcriptomic profile between those samples. Since the cells subject to fixation were harvested from the same culture as the non-fixed cells, biological variation between the two datasets is expected to be small. If the methanol fixation indeed does not result in any significant changes to the RNA profile, then the correlation between the live and fixed transcriptomic datasets should be high, and comparable to within-dataset correlations. To validate this hypothesis, we first randomly selected three cells from each of the live and fixed datasets and made scatter plots to visualize the pairwise similarity between single cells at the gene level (Figure 1C). Indeed, scatter plots look as expected, with high expression genes between single cells correlating closely while low expression genes are more broadly dispersed, with generally good correlation across all genes (24). We also calculated Pearson correlation coefficients for each pair. As expected, the r^2^ values are consistently high for both cells compared within live or fixed datasets, and between live and fixed datasets. These r^2^ values are also comparable to those found in other published single-cell cross-correlation analyses (25). To further confirm these results, we then calculated the pairwise correlation for all the cells we profiled, visualizing the results in a heatmap (Figure 1D). Overall, the correlation between all cells is high, between 0.7 and 0.9. The annotation bar indicates the label of each cell, live or fixed, and the intermixing of labels indicates that the degree of correlation is not clustered by sample type, suggesting that the methanol fixed cells do not show a major difference from the live cells. These results show preliminarily that methanol fixation does not result in any obvious changes to the transcriptomic profile of single cells.

### Methanol fixation does not affect cell-type identification, clustering, and biological inference

We found that methanol fixation does not dramatically change single-cell RNA transcriptomic profiles, but scRNA-seq is most commonly used to perform cell-type identification and clustering, therefore we further explored our data using classification methods to ensure fixation does not affect these types of analyses and downstream biological inferences. Principal component analysis (PCA) is a commonly used technique in single-cell RNA-seq analysis (26). It identifies the coordinate system that represents the greatest variance in the data, and projecting data points in this new coordinate system, thus is able to visualize the differences between groups of data points and cluster similar data points together. To see whether single cells could be grouped by their fixation treatment, which would indicate that there is the variance between the two treatment groups, we applied PCA on our data and checked the first several principal components (PCs) for separation between groups of cells. We found that the top three PCs show meaningful separations (Figure 2A): The first PC, which represents the greatest degree of variance, separates cells according to their cell type; the second PC appears to correlate with the cell cycle phase of each cell, and after normalizing for cell cycle effects, we observe that cells become clustered by their treatment condition (Figure 2A middle and bottom rows). This suggests that among all the factors for cell classification, inherent differences in cell type remain the most prominent, and when performing cell clustering analysis, any significant biological differences between cell types are unlikely to be obscured by the effects caused by fixation.

**Figure 2.**
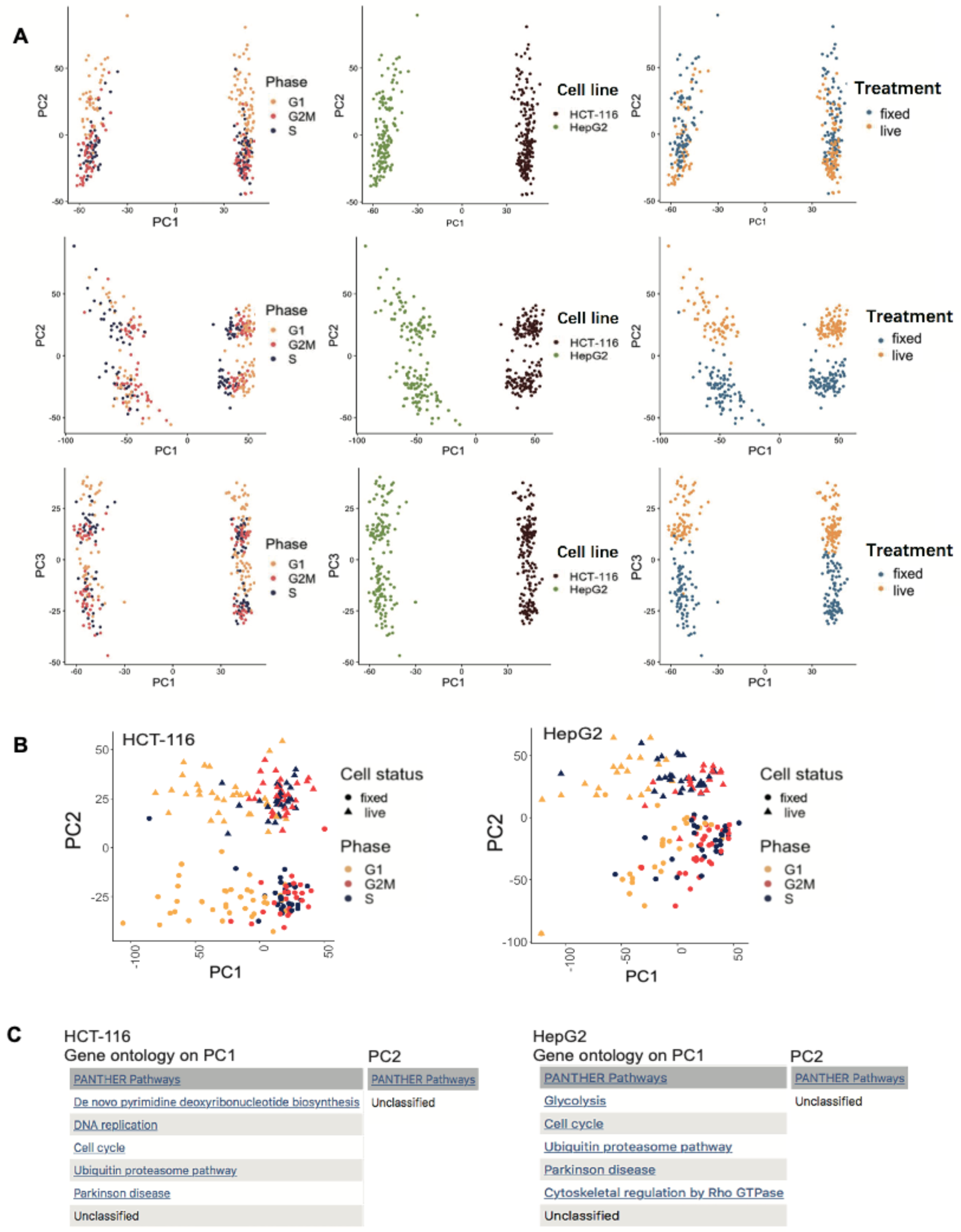
Principal component analysis of data generated from two cell lines. **(A)** PCA visualizing different treatments and annotations. The first row visualizes PC1 and PC2. The third row visualizes using PC1 and PC3. The second row visualizes PC1 and PC2 after cell cycle effect removal. Cells in the same column are annotated using the same terms. Cell type confers the greatest degree of variance in the dataset as shown by the first PC, followed by cycle and fixation effect. Key biological differences between cell types are not obscured by the fixation effect. **(B)** PCA of the individual cell line. Both PC1s are separated by cell cycle effect, while PC2s are separated by the fixation treatment. **(C)** Gene ontology terms of 500 genes with the top contribution in separating the first and second PCs in both cell lines. We further validated the smear pattern in Figure 2A was caused by cell cycle effect and the separation between live and fixed cells is not caused by biological reasons.

To determine the specific genes and possible pathways that are responsible for the separation between live and fixed cells, we performed PCA on each cell type separately, and as expected in this analysis PC1 showed separation between cells according to cell cycle phase whereas PC2 was by treatment conditions (Figure2B). We then extracted the top 500 highly variable genes from PC1 and PC2 in each cell line, and performed Gene Ontology (GO) Analysis (27) on these genes (Figure 2C)(Supplementary Table S1). High contribution genes from PC1 correspond to biological pathways involved in cell cycle processes and control for both cell types analysed, which is expected based on our previous analysis. Genes that are heavily loaded in PC2, which separate the cells by their fixation treatment, did not correspond to any known biological pathways in GO. This result suggests that the separation between live and fixed cells is likely not regulated by any specific biological mechanisms, but rather by technical factors.

### Genes that drive live and fixed separation show greater variation in expression level

To explore the PCs with the strongest variation in more detail, we studied the statistical features of the top 500 loading genes in PC1 and PC2. Two sets of genes from both PCs were extracted and their relative expression abundances were studied. Specifically looking at those genes with high loading in PC2 that are responsible for the separation of live and fixed groups in this PC, we compared their average expression between live and fixed cells and found that the key difference is that low-expression genes are generally less detected or less expressed in the fixed cells (Figure 3A). We do not observe this phenomenon with genes from PC1 (cell cycle), indicating that this is unlikely to be caused by any technical limit of detection (LOD) – a compromised LOD would affect all low-expression genes in the sample and therefore would appear in both PCs, which is not the case. In addition to the changes to the mean expression level of low-expression genes, we also observe differences in the variability of the gene expression level when comparing the genes from the two different PCs (Figure 3B). The coefficient of variation (CV) across cells of the gene expression level for genes contributing to PC1 (cell cycle) is comparable between the fixed and live groups, suggesting that cell-cycle related genes are detected with similar consistency in each cell population regardless of the treatment condition. Genes contributing to PC2 (fixation effect), however, show notably higher variation in fixed cells than in live cells (Figure 3B bottom panel). These results suggest that the effect of methanol fixation could be specific to those genes. To summarize this part, methanol fixation does not cause consistent signal lost for the whole transcriptome, but rather stochastically across all cells for genes involved in PC2 separation. Therefore, genes separating PC2 may share common features that make them specifically affected once fixed. In conclusion, the discrepancy between live and fixed cells is likely not due to any biological process of the cell that is induced by methanol treatment.

**Figure 3.**
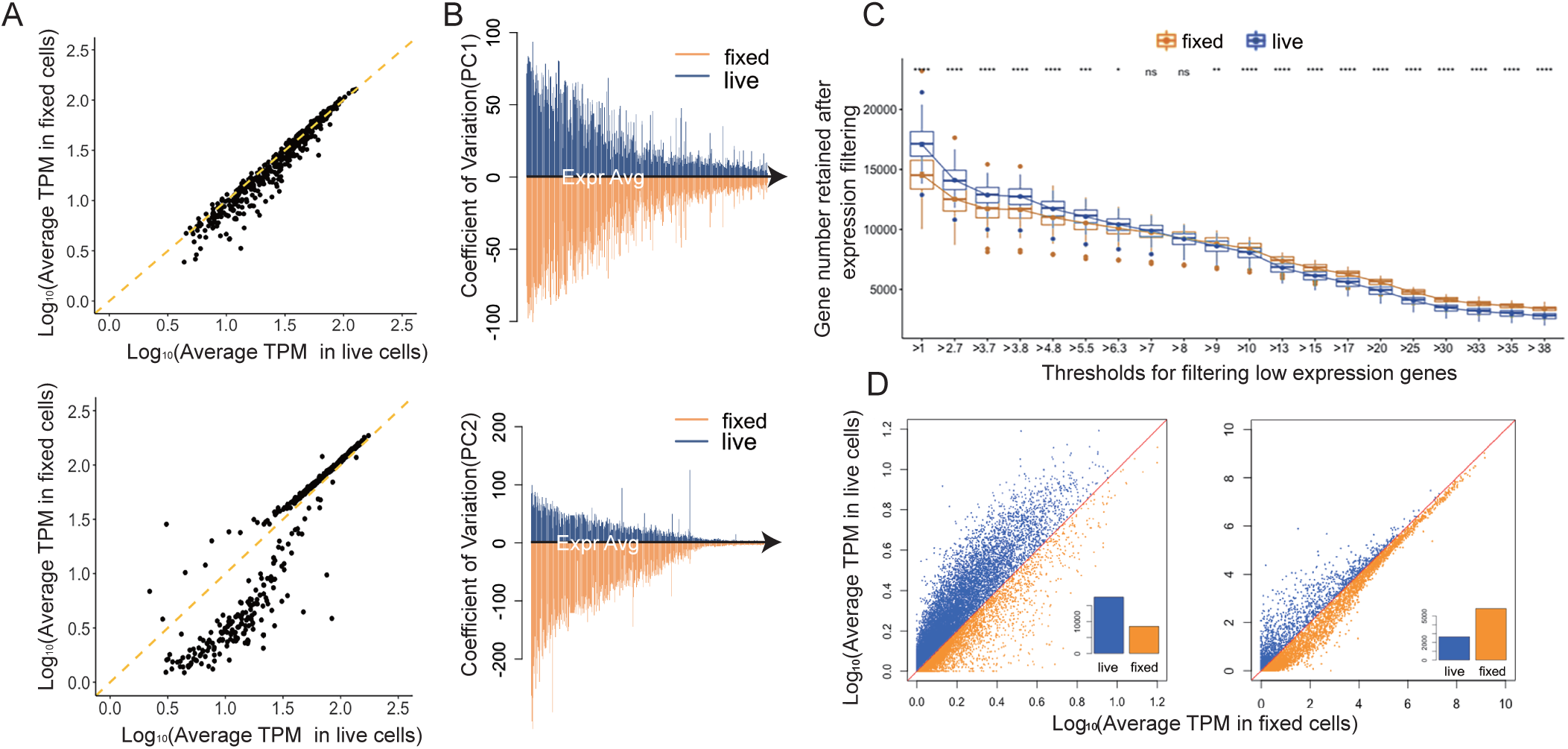
Differences in statistical features of genes with the top contribution in driving variation between live and fixed cells. **(A)** Comparison of relative expression of 500 genes with the top contribution in PC1 and PC2 between live and fixed cells. Expression of PC1 genes correlated well while in PC2 the trend was incoherent for genes with different expressions level, which indicates genes heavily loaded in PC2 may be responsible for the separation between two groups of cells. **(B)** Comparison of expression variation of genes with top contribution from PC1 and PC2. In the top panels of Figure 3B, we take genes that are heavily loaded in PC1 respect for live cells and fixed cells. Then, we computed the coefficient of variation (CV) of each gene across all cells. The CVs for each gene are then plotted against that gene’s mean expression level, separately for live (blue) and fixed (orange) cells. Genes with the top contribution in PC2 holds much higher variation compared with PC1 genes. **(C)** Comparison of gene detection number after expression filtering. A series of thresholds were set up for different sensitivity requirements. The detection number in fixed cells gradually surpass live cells once the threshold increased (nsP>0.05, *P<0.05,**P<0.01,****P<0.0001). **(D)** Relative abundances of genes with high (>30 TPM) or low (<5TPM) expression, the inset bar charts compare the quantities of genes which have higher expression in either live (blue) and fixed (orange) cells. For low expression genes, they are generally more abundant in live cells. Genes with higher expression are more abundant in fixed cells.

Since scRNA-seq is known to suffer from dropout events in gene detection, we wondered if fixation exaggerates this phenomenon. To better evaluate the dropout frequency over the entire transcriptome, we set a series of increasing gene expression level thresholds for defining detected genes. For each threshold, we used boxplots to visualize the number of genes with expression levels greater than this threshold (Figure 3C). As expected, when the threshold for gene filtering is low, live cells have more genes detected overall; but somewhat surprisingly, as the gene expression threshold gradually increases, a greater number of genes is detected in fixed cells. This result shows that fixed cells tend to have more dropout events for low expression genes, but retain higher expression genes more robustly. We further illustrate this by extracting genes with either high or low expressions (gene expression (TPM) >30 high or <5 low), and for each group, visualizing the relative correlation between the mean expression level for each gene (Figure 3D). The result shows low expression genes are more abundant in the live group than the fixed. The inset graph shows the quantitative comparison of gene numbers above or below the diagonal line. The trend was reversed for highly expressed genes that their expressions are more abundant in fixed cells. Based on these results, we concluded that the frequency of dropout and the relative quantitative expression are different between live and fixed cells. And the methanol treatment differentially affects genes with different expression levels.

### Longer and higher GC transcripts are more severely affected by fixation

We sought to find features that are shared among those genes that are most affected, however, features other than abundance can only be described for transcripts, not genes. Abundance measurements at the gene level represent the contribution from multiple transcripts, potentially of widely varying lengths and sequence properties. Therefore, subsequent analyses used transcript level abundances to shed light on potential molecular features or mechanisms that lead to certain types of transcript molecules being affected more by methanol fixation.

Length and GC content are two important features to be considered. To visualize the GC and length level of specific transcripts against the rest of transcriptome, we sorted all transcripts by their length and GC content and made rank-order plots. In these plots, each dot can be located by a gene’s feature and its corresponding rank, in the increasing order. In the GC content plot, we highlighted top contribution genes from PC1 (cell cycle) and PC2 (fixation effect) using coloured dots, while remaining transcripts are plotted in grey (Figure 4A). Compared with PC2, PC1 genes have more even distribution along with the line plot compared to those from PC2. Most PC2 transcripts are restricted to the higher GC content part, which indicates that transcripts separating fixed cells from live ones have higher GC base-pairs in the sequence in general. A similar pattern was revealed when the same analysis was done for transcript length (Figure 4B). To compare the length and GC content of transcripts from both groups, p-value was calculated for each using T-test, and a statistically significant difference was found between genes contributing to PC1 and those contributing to PC2 (Figure 4C). The fixation effect is more prominent for long and high GC transcripts, which are features of transcripts that are causing non-biological separation between live and fixed cells.

**Figure 4.**
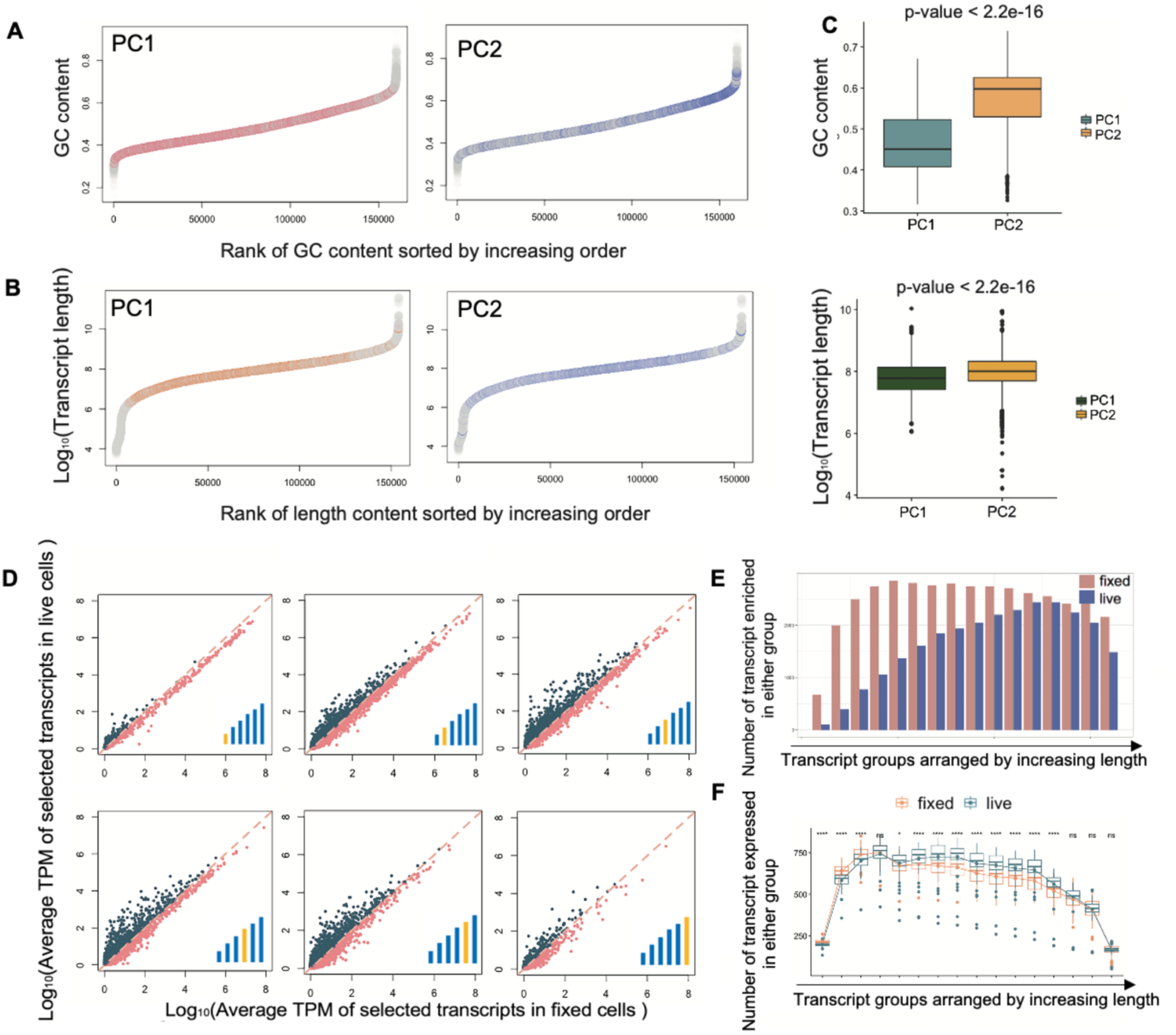
Molecular features of transcripts separating PC1 and PC2. **(A)** Plots of GC content and corresponding rank for the whole transcriptome. Highlighted events are those with top contributions in PC1 (left) and PC2 (right). GC contents of PC2 transcripts are generally higher compared with PC1 transcripts. **(B)** Plots of length and corresponding rank for the whole transcriptome. Highlighted events are those with top contributions in PC1 (left) and PC2 (right). Lengths of PC2 transcripts are generally higher compared with PC1 transcripts. **(C)** Comparisons of GC (top) and length (bottom) ranking of transcripts with top contributions in PC1 and PC2. P-values show differences between live and fixed groups are both significant. Transcripts with top loading in PC2 are generally with longer lengths and higher GC contents compared with those in PC1. **(D)** Comparisons of relative abundances of transcripts with different lengths. We put a set of bars with increasing height at the bottom right corner to represent the transcript lengths. Highlighted bars represent the relative length of transcripts employed in that plot. As transcripts get longer, they gradually become more abundant in live cells than fixed cells. **(E)** Comparison of abundant transcript quantity in live and fixed cells. Groups separated by length. **(F)** Comparison of transcripts detection number. Groups are separated and arranged by increasing length. The number of transcript detection varies as length changes. Statistical significance p-values are determined by t-test and indicated with asterisks (nsP>0.05, *P<0.05, ****P<0.0001).

To visualize how transcript features correspond with the fixation effect an individual receives, we compared relative expression level and transcript detection number. For abundance comparison, we separated transcripts into 16 groups with equal size according to length (6 plots with increasing order of length were selected) (Figure 4D, Supplementary Figure S6-S7). We compared relative expression by correlation plot, and the comparison pattern differs as transcript length varies. Then for each group (16 in total), we counted transcript number above or below the diagonal line, which stands for if a transcript holds higher expression in live or fixed cells, to compare the number of transcripts that are enriched in either group (Figure 4E). The gradually changing trend illustrates that shorter transcripts are more enriched in the fixed group, yet longer transcripts have more equal expressions for both groups. The transcripts detection number shows similar variations (Figure 4F). For shorter fragments, more transcripts are detected in fixed cells, whereas longer fragments are detected more frequently in live cells. These analyses suggest that transcripts receive different degrees of effects result from different lengths and GC contents. And transcripts with longer lengths and higher GC contents are more likely to be affected by methanol fixation.

### Transcripts that are both long and GC-rich are the most affected by fixation

So far, fixation is shown to differentially affect transcripts with different GC content and length. However, how these features result in the discrepancy is still unclear. Since the transcript quantification is calculated from aligned read count, which cannot reflect the completeness of sequenced fraction per transcript, we next assessed the read coverage on the transcriptome. Since Smart-seq2 is based on template switching by poly-A tail selection, fewer intact mRNA templates will result in higher 3’ end coverage, then we performed mapping coverage analysis to see if there is any alternation in coverage pattern. Transcripts are separated into 10 groups with equal size based on length. For each group, we overlapped the coverage pattern of live and fixed cells and compare the difference. The coverage traces are mostly the same for shorter transcripts (Figure 5A). However, as transcripts get longer, more reads are stacked at the 3’ prime in fixed cells. The degree of discrepancy is enlarged by increasing fragment size.

**Figure 5.**
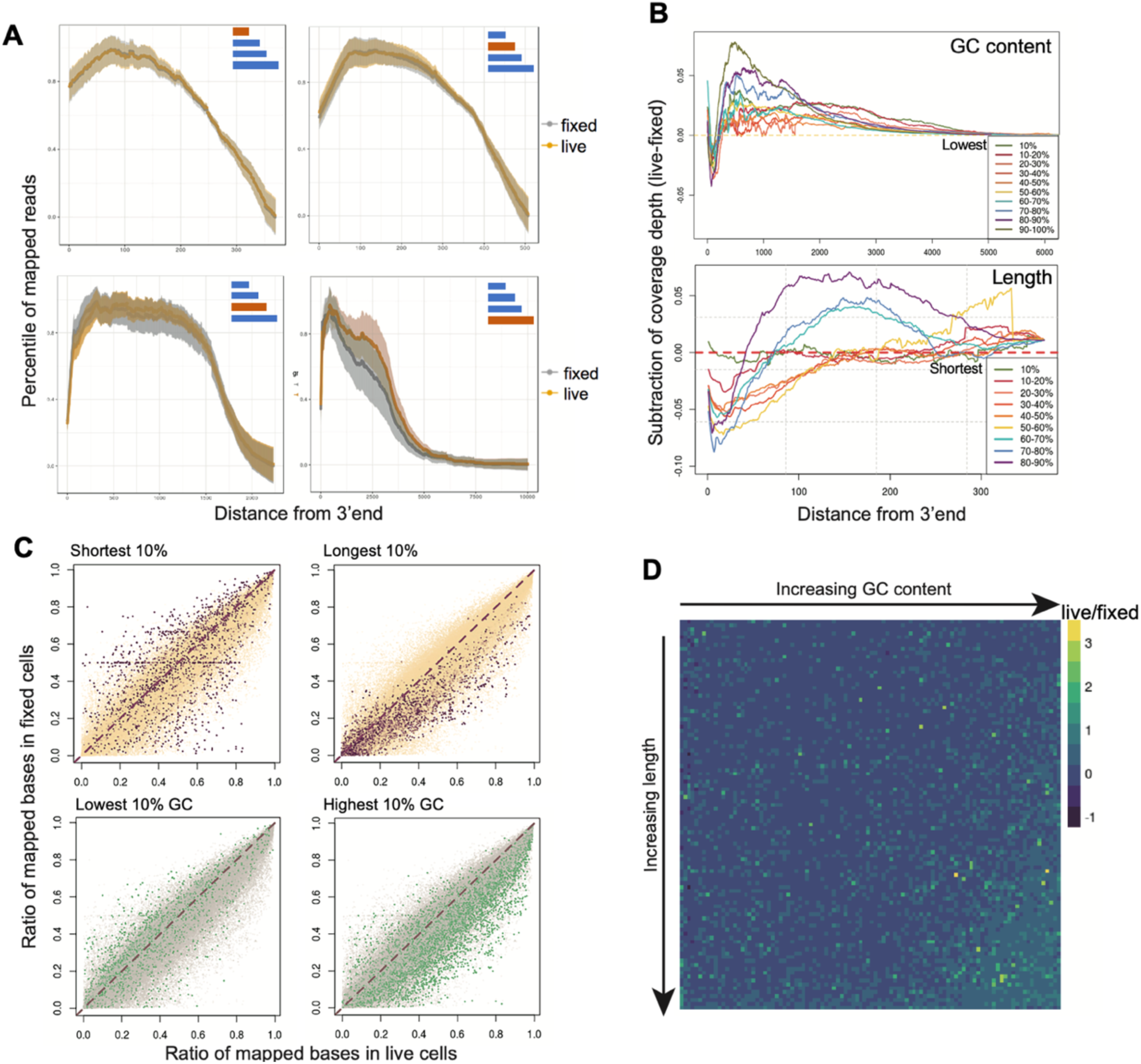
Comparison of mapping features between live and fixed cells. **(A)** Mapping coverage of transcripts grouped by different lengths was used to show the variance in transcript mapping depth between live and fixed cells. Highlighted bars in the top-right corner shows the length of transcripts involved in that plot. Bias at 3’ end in fixed cells is more obvious for longer transcripts. **(B)** Difference in the mapping depth between groups. Ten groups of transcripts were separated by either GC-content (top) and length (bottom). The difference in depth is plotted against distance from 3’end to show how the variance changes the length of each transcript. Both plots show the difference between mapping coverage is more notable for transcripts which are GC-rich or long. **(C)** The mapping ratios for each transcript were compared using coverage integrity correlation. Transcripts with top or bottom 10% rank in length and GC content are highlighted in each correlation plot. The coverage integrities of both long and GC-rich transcripts are higher in the live group than the fixed group. **(D)** Visualization of the ratio of live/fixed mapping integrity showing transcripts with better coverage. Transcripts are sorted and grouped by length and GC content; each unit represents an average ratio for those transcripts. In the corner containing transcripts with longer and GC-rich transcripts, live cells are shown to have more complete mapping compared with fixed.

Next, we directly compared the degree of coverage deviation in groups with different lengths or GC contents and studied how the variance changes along with the transcriptome from 3’ to 5’ end. We plotted curves representing the difference of mapping depth along with the transcripts and merged them into one, with length scale (Figure5B). We observed longer transcripts are shown to have more discrepancy in coverage pattern once fixed (Figure 5B, top panel). The same effect is observed for GC content (Figure 5B, bottom panel). According to the analysis above, we found that except for extremely long transcripts, fixation didn’t introduce strong 3’ bias which possibly indicates RNA degradation. In conclusion, the coverage patterns of transcripts are affected by fixation, the degree of effect increases for transcripts which are longer and higher in GC.

Since the transcript coverage is the aggregation of concatenated reads, calculated based on the whole transcriptome, it lacks the information of the mapping percentile for each transcript. To better quantify the mapping completeness, we counted how many bases are mapped for individual transcript and calculated a mapping ratio ranging from 0 to 1 and plotted the ratios (Figure 5C). By highlighting the top 10% longest and shortest transcripts, the position of selected events is shown. The shortest transcripts have similar degrees of correlation and the overall pattern was almost symmetrical. Longer transcripts are quite skewed towards the live axis, showing higher mapping integrity in the live group. The GC content plot also shows that high GC content transcripts have higher mapping percentile in live cells, while low GC content transcripts seem to have higher mapping percentile in fixed cells. To determine if these two factors co-occur, we get a quotient of mapping percentile for each transcript from live and fixed groups and binned those numbers into 10000 groups by 100×100 combinations of transcript length and GC content. We plotted a heatmap using a relative mapping ratio, in which each axis is arranged by increasing length or GC content (Figure 5D). The corner representing transcripts that are both long and are high in GC content shows higher mapping percentiles in live cells, which indicates that the effect of methanol fixation effect is more severe for these transcripts that are both long and GC-rich.

Although the fixation effect can be revealed in PCA clustering, it is presented as the second largest variation. Since we concluded that the fixation separation was mainly caused by long and GC-rich transcripts, we then explored whether the fixation effect exhibited in the first PC could be revealed by only selecting those long or GC-rich transcripts. To do so, we binned all transcripts by their GC-content into five bins with increasing GC-richness; we also binned all transcripts by their length into five bins with increasing length. We then performed PCA using increasingly higher threshold cut-off for both GC and length, meaning the set of transcripts used for performing PCA were increasingly restricted to those that are long and have very high GC content. As the threshold for GC and length increased, the amount of variation between live and fixed groups that is explained by PC1 also increased (Figure 6A), compared to when all transcripts are included, PC1 predominantly shows variation arising from differences in the cell cycle. This is presumably due to the increased weight of transcripts with long length and high GC content that contributes to the separation of cells caused by fixation. To show this more clearly, from each PCA performed using different transcript length-GC selection thresholds, we plot PC1 loadings annotated by treatment condition, in which we observed that although the magnitude of separation is reduced when fewer high length-GC transcripts were used, the separation of fixed and live cells becomes less ambiguous in PC1 (Figure 6B). We then identified the top 100 most highly loaded transcripts in PC1s and visualized their length and GC content. As the threshold for GC and length increased, the length and GC content of the top loaded transcripts also gradually increased (Figure 6C). This series of analyses illustrate that longer and higher GC transcripts indeed separate cells based on fixation, indicating that such types of transcript molecules are more likely to be affected by fixation during sample preparation.

**Figure 6.**
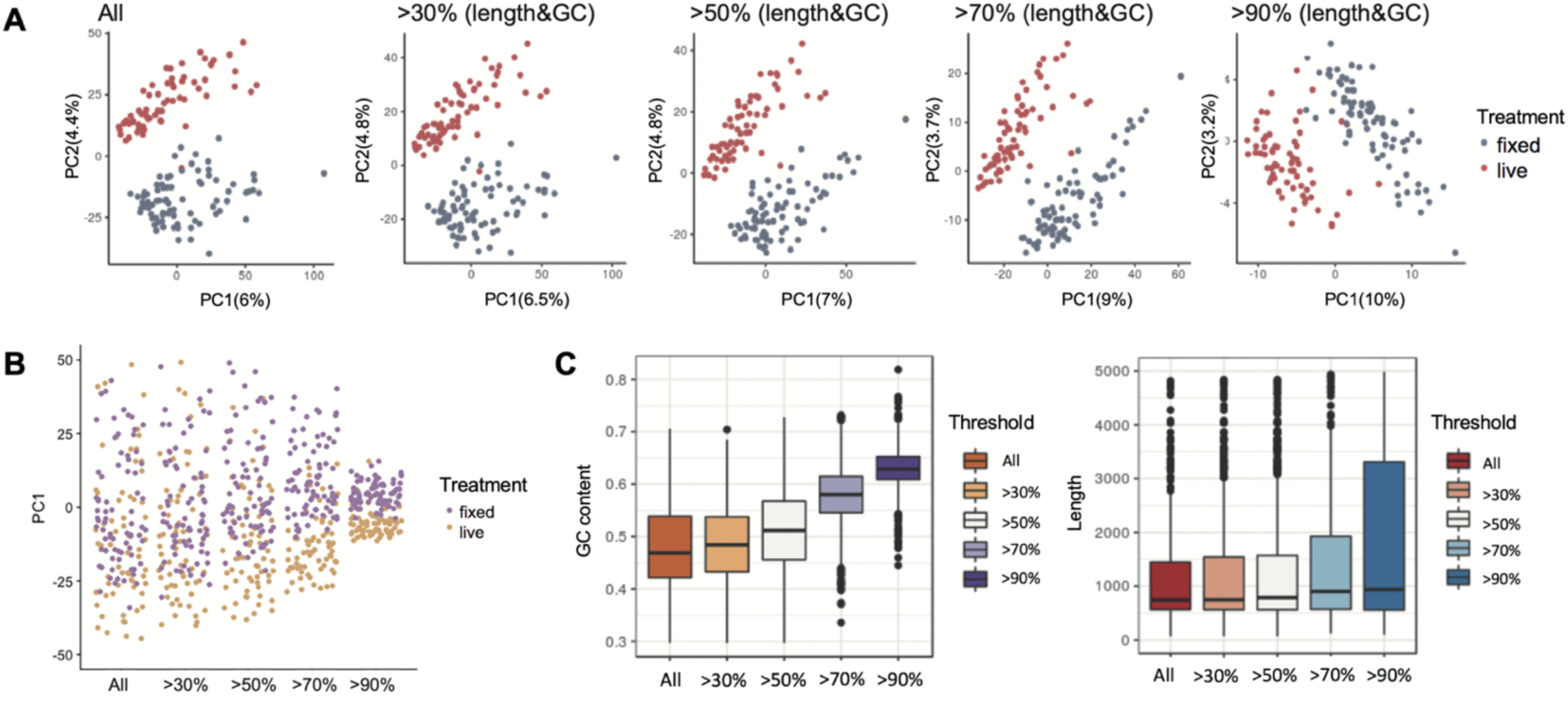
Transcripts with longer lengths and higher GC contents separate live and fixed cells. **(A)** PCA performed using different transcripts sets. In each plot, transcripts are selected based on lengths and GC contents thresholds. Transcripts selected were used for analysis and plotting. As transcripts with longer lengths and higher GC are used for PCA, PC1 is gradually dominated by the fixation effect. **(B)** PC1 loadings of cells in PCAs performed with different transcripts sets. Although the separation force increases in general, the distinction between the live cells and fixed is gradually clear. **(C)** With PCAs performed with increasing lengths and GC contents thresholds, corresponding length or GC content statistic of the top 500 transcripts from PC1s was plotted. This result validates the gradually exposed fixation effect is essentially caused by transcripts that are long and GC-rich.

### Methanol fixation renders less effect on the sequencing result generated from 3’bias method

Since Smart-seq2 produces full-length cDNA libraries, the uneven influence received in different transcripts can be observed in mapping coverage and gene quantification. On the other hand, protocols preserving only the 3’ end of cDNA library such as Drop-seq may be less affected by methanol fixation (28), since only one end of the transcript is counted during quantification and analysis. Therefore, even if the RT and amplification are hindered by changed mRNA structure, the fixation effect will be less significant than that seen in Smart-seq2 data.

To validate our hypothesis, we first simulated Drop-seq data by mapping reads generated in Smart-seq2 to the 3’ end of transcriptome reference. By doing so we can produce a dataset resembling the features of Drop-seq since both methods are based on template switching strategy but Drop-seq only preserving 3’ end of the library. In the PCA plotted with 3’ end mapped data, although we still observe the distinction between cells processed with different treatments, the separation is much less clear (Figure7A), and the two clusters appear merged into one. Compared with full-length Smart-seq2 data, the simulated 3’ biased data shows little fixation effect. To examine if the merging of two clusters is caused by different alignment procedures, we also mapped original reads to the 5’ end of the transcriptome. When performing PCA with 5’ end mapped data, the distance between two clusters enlarged and the separation is distinct (Figure 7B). The result shows data similarity between live and fixed cells differs between 3’ and 5’ end of the transcriptome.

**Figure 7.**
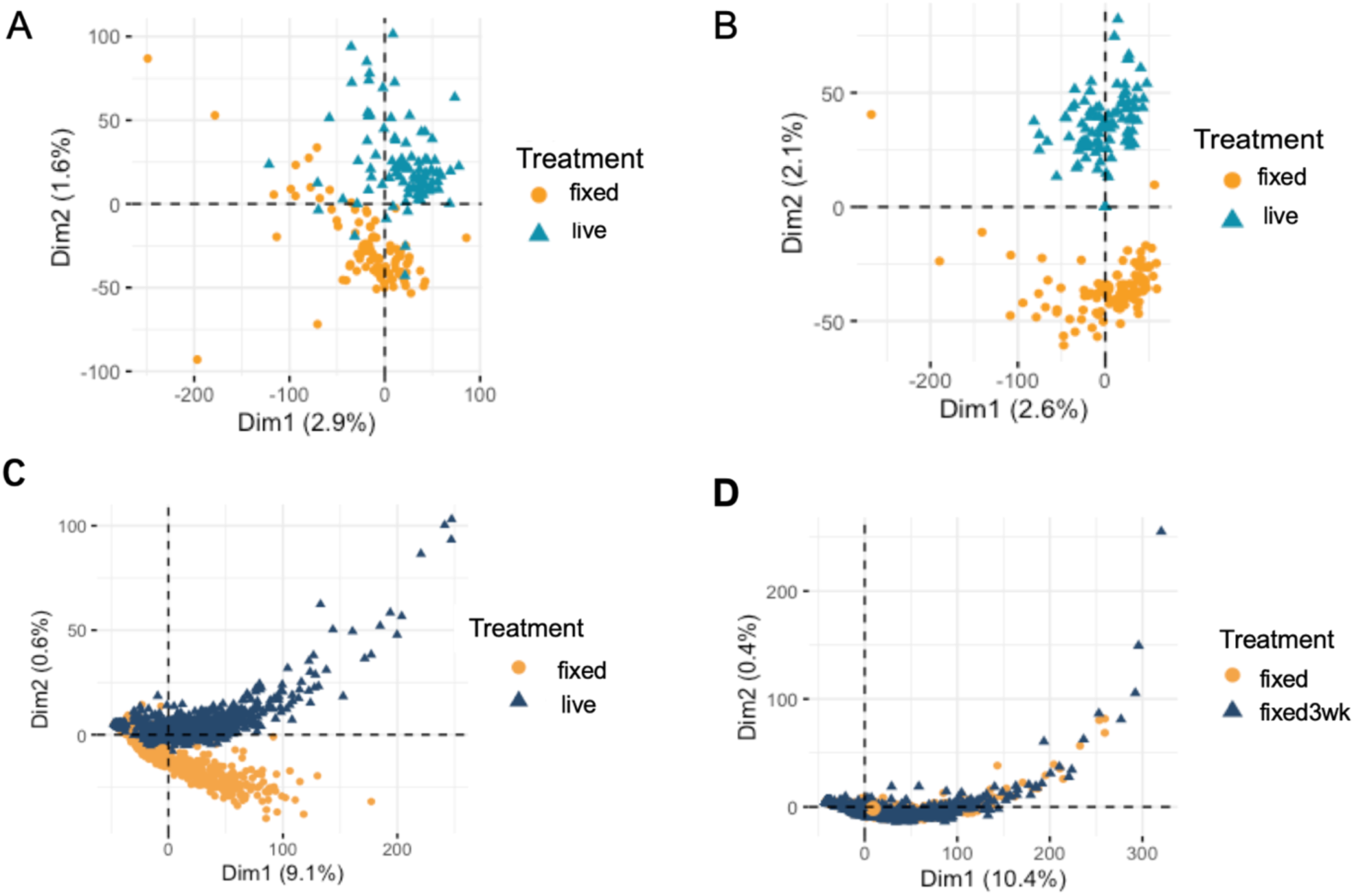
Analysis of the fixation effect in 3’ end biased sequencing data. (A) PCA clustering using smartseq2 data counting only 3’ end to simulate Drop-seq data. While the separation still exists between live and fixed cells, two clusters are not totally separate from each other. (B) PCA clustering using data counting only 5’ end of the transcriptome. The separation between live and fixed cells is clear. (C) PCA clustering including live and fixed cells generated from Drop-seq. Cells from two types are partially merged without strong separation. (D) PCA including fixed and fixed cells with 3 weeks storage. Fixed cells with different storage conditions are clustered together.

We also used published data to explore the fixation effect in a real Drop-seq experiment (14). Three sets of data generated from the HEK cell line were analysed, which included live cells, fixed cells, fixed cells with three-week storage. In the PCA including live and fixed cells, although the separation still exists, two clusters are partially merged, which indicates the fixation effect is much weaker in Drop-seq compared with Smart-seq2 (Figure 7C). When we performed PCA using cell groups that are both fixed but with different storage duration, we saw once cells are fixed before sequencing, the similarity of their transcriptomes is high. This result indicates fixation effects are consistent and not affected by the cell storage time.

This result validates our hypothesis that fixation can affect data differently according to the methods used for library construction. For protocols utilizing template switching strategy, 3’ end data are less affected by fixation compared with full-length data.

## Discussion

To elucidate the effect of methanol fixation on single-cell RNA-seq data, we performed a series of comparative analyses and proposed a potential mechanism for how the fixation effect occurs. For all comparisons carried out at the gene level, the results show that fixation data is capable of revealing biological insights that are typically sought in scRNA-seq experiments. Although subtle discrepancies were observed in part of our analysis, they did not obscure the key biological features of cells and key biological differences between cell types. We further investigated these effects at transcript-level to hone in on the source of the observed differences, using both expression abundance data and raw sequencing data to uncover more in-depth insights to account for the observed discrepancies. Using transcript-level information, we showed that length and GC are key properties that correlate with the degree of fixation effect each transcript receives. Specifically, longer fragments with higher GC content are shown to be more affected by fixation in quantification and mapping integrity. Based on structural considerations, we hypothesize that transcripts that are long and high in GC are more likely to have complex higher-order structures, which makes them more difficult to fully recover from methanol fixation even after rehydration (29). Unlike mRNA with simple structures, those with more complex structures may be altered and hinder downstream reactions such as RT and amplification. An important insight from our results is that for users of scRNA-seq who wish to investigate differences in splice isoforms, live samples will be more reliable since the full-length information of transcriptome is better preserved; methanol fixation will result in skewed abundance readouts from those transcripts with high GC and long length.

In addition, fixation effects are more obvious in Smart-seq2 data compared with Drop-seq. In both simulated and real Drop-seq data (Figure 7A,7C), we observed less separation between live and fixed cells compared with Smart-seq2, therefore illustrating that the fixation effect is not observable in 3’ end sequencing. It is possible that since the oligo-dT primers bind to the poly-A tail at the 3’ end to initiate the RT, the integrity of the 3’ end is more likely to be protected and captured, whereas the subsequent template switching step that occurs at the 5’ end is more likely to be affected. An inefficient template switch then leads to incomplete DNA elongation and finally affecting data quantification in fixed cells. Based on our observations and mechanistic hypothesis, methanol fixation influences the data by introducing barriers to RT, which will occur more often in transcripts with complex secondary structure, therefore the fixation effect will not be observed if sequencing only one end of the transcript.

Besides the loss of mRNA with more complex secondary structures, we also observed a general dropout trend of low expression genes. By checking the sequence features of the corresponding transcripts, we found the low expression is not related to GC content and length. Instead, this may indicate the dropout detection of low expression genes is exaggerated by fixation. For research focusing on transcripts with low abundance, methanol fixation can lead to reduced or even complete loss of the target. For all the fixation effects described above, they are recognized as fixation complications, without affecting overall interpretation and analysis of biological processes. We also found that although the discrepancy between live and fixed sample is not primarily caused by RNA degradation, it is crucial to include protective reagents such as RNase inhibitors to all buffers and reagents during the methanol fixation and rehydration steps to prevent degradation – something that was not previously highlighted or emphasized in existing fixation protocols.

In addition, fixation induced expression changes are also seen when comparing relative abundance between live and fixed cells (eg. Figure 3D, Figure 4D). When using Smart-seq2 data for such analysis, some transcripts show higher expression in fixed cells rather than in live cells. This begs the question of whether some genes are enriched due to fixation. Although the elevated expression of partial mRNA in fixed cells in some analysis seems contradicted, this phenomenon can be explained from the perspective of library construction and data normalization. During sequencing library construction and single-cell library pooling, the protocol aims to collect equal amounts of products from individual samples. Therefore, even though there are differences in cDNA yield after pre-amplification, it can be removed by sequencing library preparation since the equal quantity of cDNA were used for downstream processing. Moreover, after data normalization, eg TPM and CPM, the resulting library size will be equalized for individual cells. In fixed cells, transcripts with inherent low expression or those with complicated structure have more dropout during library preparation, therefore loss of those transcripts makes space for remaining transcripts, which is presented as a higher expression level in fixed cells.

Our work determines the feasibility level of methanol fixation in different usage scenarios such as basic scRNA-seq compared with those focusing on transcript isoforms level studies and informs users of the types of biases that occur with methanol fixation in different experimental scenarios. Knowing how fixation affects the RNA-seq data is beneficial for researchers to make reasonable and appropriate experimental plans using methanol fixation.

## Materials and methods

### Cell line preparation and fixation

HCT-116 and HepG2 cells were cultured with Dulbecco’s Modified Eagle’s medium (DMEM) (Thermo Fisher Scientific, cat# 12100046) supplemented with 1% Penicillin-Streptomycin (Thermo Fisher Scientific, cat# 15070063) and 10% Fetal Bovine Serum (Thermo Fisher Scientific, cat# 16000044). We harvested cells at 70-80% confluence, dissociated for 2 min at 37C° using 0.25% trypsin-EDTA (Invitrogen, cat# 25200072), and quenched with growth medium. To prepare cells for different treatments, the cell suspension was separated into two parts with equal volume. We used 300gX5min centrifugation to wash the cell. The supernatant was removed, and the cell pellet was resuspended using phosphate-buffered saline (PBS) (Invitrogen, cat# 10010023). Resuspension volume was chosen to make cell concentration around 1×10^6^ to 10^7^ cell/mL. For cells to be prepared for live library generation, they were ready for immediate further processing. The rest part of the cells was kept for fixation.

Fixation and rehydration steps were performed following the protocol in (14). Ice cold 20% PBS was added to resuspend the cell pellet and 80% pre-chilled methanol was added dropwise, total volume was calculated to achieve the concentration around 1×10^6^ to 10^7^ cell/mL. After mixing PBS and methanol by gently pipetting up and down, the tube with the fixed cells was placed on ice for 20 min and transferred to −80C° for longer storage. Fixed samples were kept for one week before the following steps.

### Cell rehydration, FACS sorting, library construction, and sequencing

Cells stored in 80% methanol was transferred from −80C° to ice and centrifuged at 1500g for 3 mins. We discarded supernatant and collected the cell pellet. PBS in presence with 0.01%Bovin Serum Albumin fraction V (BSA, Thermo Fisher Scientific, cat# 15260037) and 1U/ul RNase OUT (Thermo Fisher Scientific, cat# 10777019) was used for resuspending and washing the cell pellet. Fixed cells were washed with the same washing buffer twice to remove methanol thoroughly. After washing, fixed cells were kept in the washing buffer and ready for sorting. For live cell sorting, cells were stained with propidium iodide (PI, Sigma-Aldrich, car# P4864-10ML) solution at room temperature for 10 min. Both live and fixed cells were filtered through 70μm filter to prevent cell clumping.

Both live cells and fixed cells were sorted into 96 well plates containing cell lysis buffer using BD Aria™ IIIu sorter (BD Biosciences). FSC parameters were used for singlet selection. PE-Cy5 signal was used for removing dead cells. The plates with sorted cells were vortexed and spun down at 4C°. Single-cell cDNA library was constructed using Smart-seq2 protocol (16). After obtaining cDNA libraries, cDNA library size was checked using Fragment Analyzer HS NGS Fragment Kit (1-6000bp) (Agilent formerly Advanced Analytical, cat# DNF-474-1000). Concentrations were quantified by Qubit 3.0 fluorometer (Thermo Fisher Scientific, cat# Q33216). High quality cDNA libraries without notable degradation were used for sequencing library construction.

Illumina sequencing libraries were prepared using Nextera XT DNA Library Prep Kit (Illumina, cat# FC-131-1096). The concentrations of cDNA libraries were diluted to 0.1-0.3ng/ul. Tagmentation and dual index adding were done following the protocol provided by C1 Fluidigm (Fluidigm). Single-cell library was pooled with equal volumes and sequenced using Nextseq500/550 High Output Kit v2.5 (Illumina, cat# 20024906) on Nextseq500/550 sequencer (Illumina) to get around 1.5 million paired reads for each cell.

### Data processing and analysis

Raw sequencing data were demultiplexed with adapters trimmed on Basespace (Illumina). Quality of all raw *fastq.gz files was checked using Fastqc (17). Reads are mapped to ENSEMBL human reference genome GRCh38 using Kallisto (18) for quantification. The integration of single-cell data was done by tximport (19). Cells with more than 4000 genes detected were preserved for downstream processing, resulting in 151 cells for HepG2, and 183 cells for HCT-116. For data normalization, we logged the TPM result provided by Kallisto. Matrices of gene and transcript abundance were obtained by adjusting “tx2gene” parameter in tximport.

The correlation matrix was plotted with “chart.Correlation” implemented in the “PerformanceAnalytics” package. Cell-cycle related analyses were done using Seurat R package (20). We performed Principle component analysis using FactomineR package and did all visualization using either R build-in function, ggplot2, or ggpubr. We performed Gene Ontology analysis using Gene Ontology website(http://geneontology.org). The rest of the analysis was done with custom scripts.

Length and GC information of individual transcript was calculated according to GRCh38 transcriptome sequence using custom R script. For all transcript separation based on either length or GC content, the thresholds were set to make sure that all groups have the equal number of transcripts. In the analysis focusing on transcript coverage, raw reads were mapped to GRCh38 by STAR (21). To analyze mapping coverage for the individual transcript, Aligned *bam files were converted to depth files with samtools depth function in SAMtools (22), which shows the number of reads mapped to each base pair on each chromosome. Reads in depth file were then assigned to specific transcripts using bedtools intersect function in BEDTools (23). Downstream analysis and visualization were performed in R.

To map Smart-seq2 reads to the 3’ end and 5’end of the transcriptome, we first trimmed the GRCh38_RNA_latest.fna to build the reference, then we performed alignment and quantification with Kallisto. Gene expression matrices of Drop-seq data were downloaded from GEO (accession numbers GSM2359002, GSM2359003, GSM 2359005).

## Supporting information

topcontributiongenelist

## Accession number

GSE150993

## Acknowledgement

We thank our lab member Qiuyu Jing for inspiring discussion and proof-reading as well as Xuemeng Zhou and Qinghong Jiang for their help with Linux coding.

