## supplementary data for "Effect of methanol fixation on single-cell RNA sequencing data"

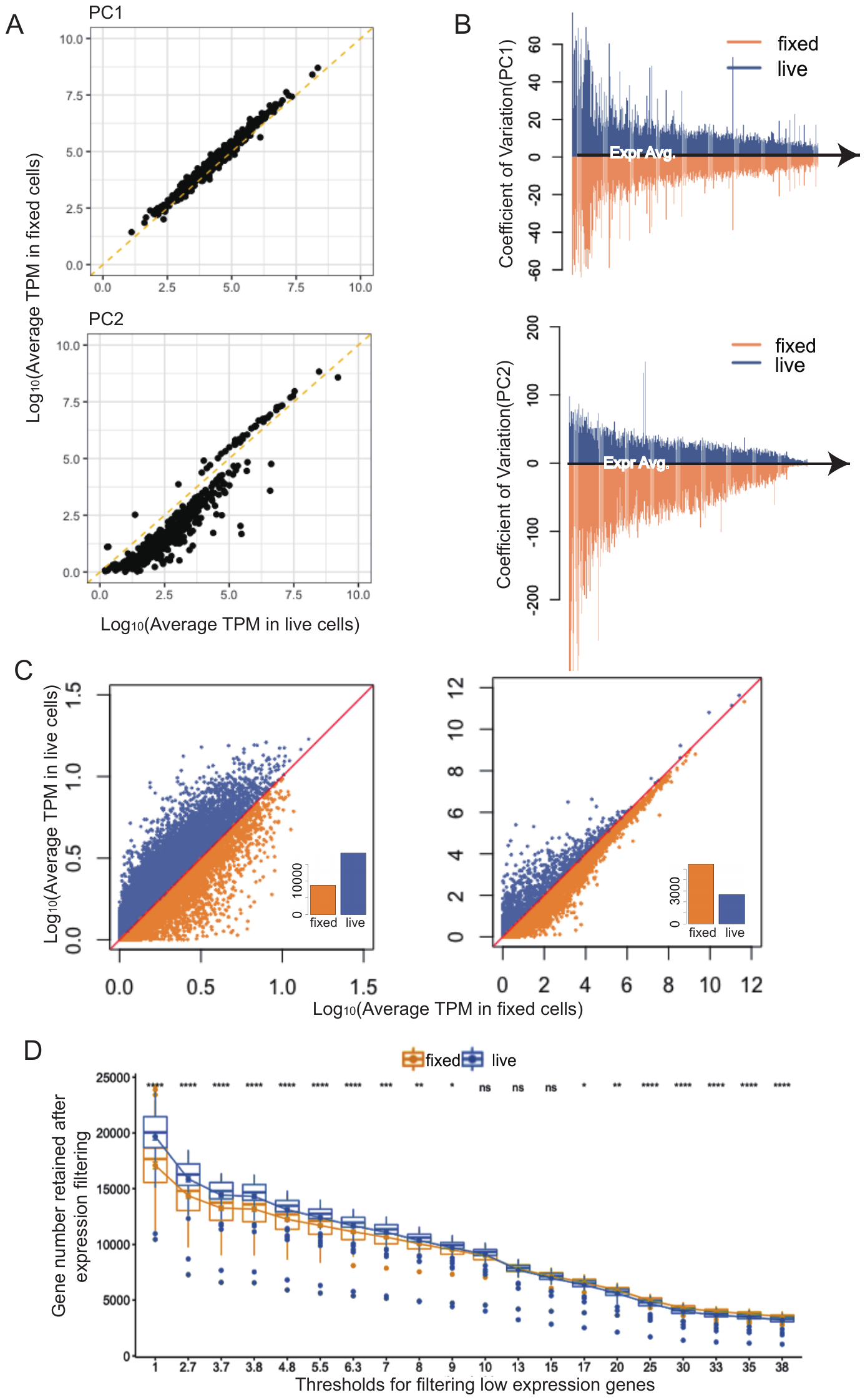

**Supplementary Figure 1.** Statistical features of genes with the top contribution for driving different PCs between live and fixed cells in HepG2.

**(A)** Comparison of relative expression of 500 genes with the top contribution in PC1(top) and PC2(bottom) between live and fixed cells.

(**B**) Comparison of expression variation of genes with top contribution from PC1(top) and PC2(bottom)

(**C**) Comparison of gene detection number after expression filtering.

(**D**) Relative abundances of genes with high (>30 TPM) or low (<5TPM) expression, the inset bar charts compare the quantities of genes that have higher expression in either live(blue) and fixed(orange) cells.

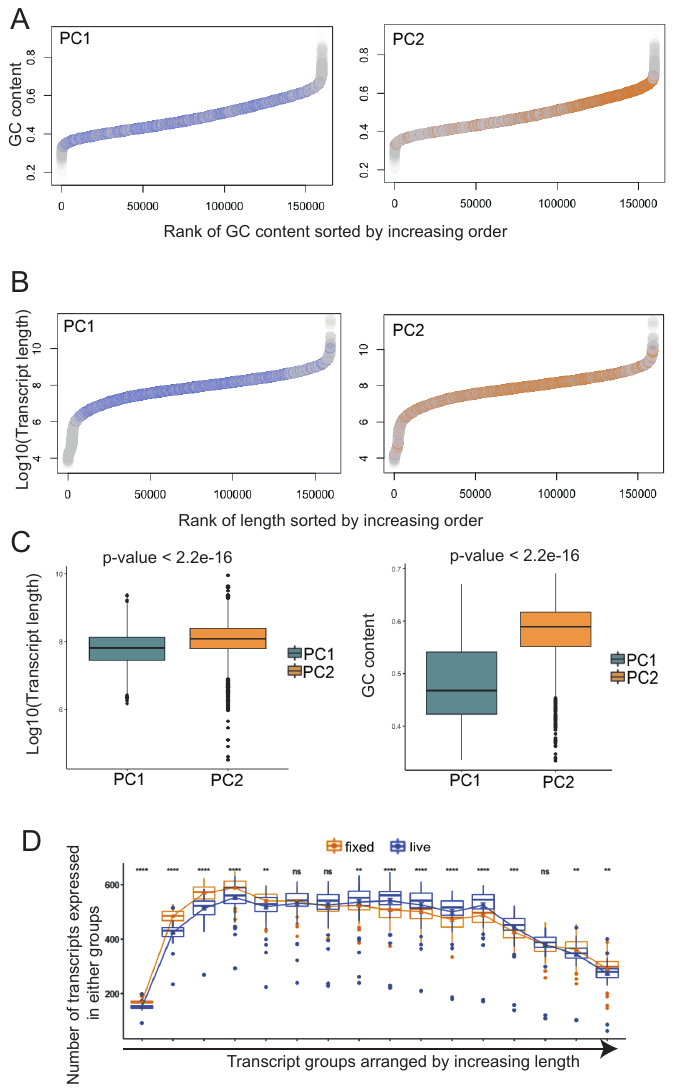

**Supplementary Figure 2.** Molecular features of transcripts separating PC1 and PC2 in HepG2.

(**A**) Plots of GC content and corresponding rank for the whole transcriptome. Highlighted events are those with top contributions in PC1 (left) and PC2 (right).

(**B**) Plots of length and corresponding rank for the whole transcriptome. Highlighted events are those with top contributions in PC1 (left) and PC2 (right).

(**C**) Comparisons of GC (top) and length (bottom) ranking of transcripts with top contributions in PC1 and PC2. P-values show differences between live and fixed groups are both significant.

(**D**) Comparison of transcripts detection number. Groups are separated and arranged by increasing length. The number of transcript detection varies as length changes. Statistical significance p-values are determined by t-test and indicated with asterisks (nsP>0.05, *P<0.05, ****P<0.0001).

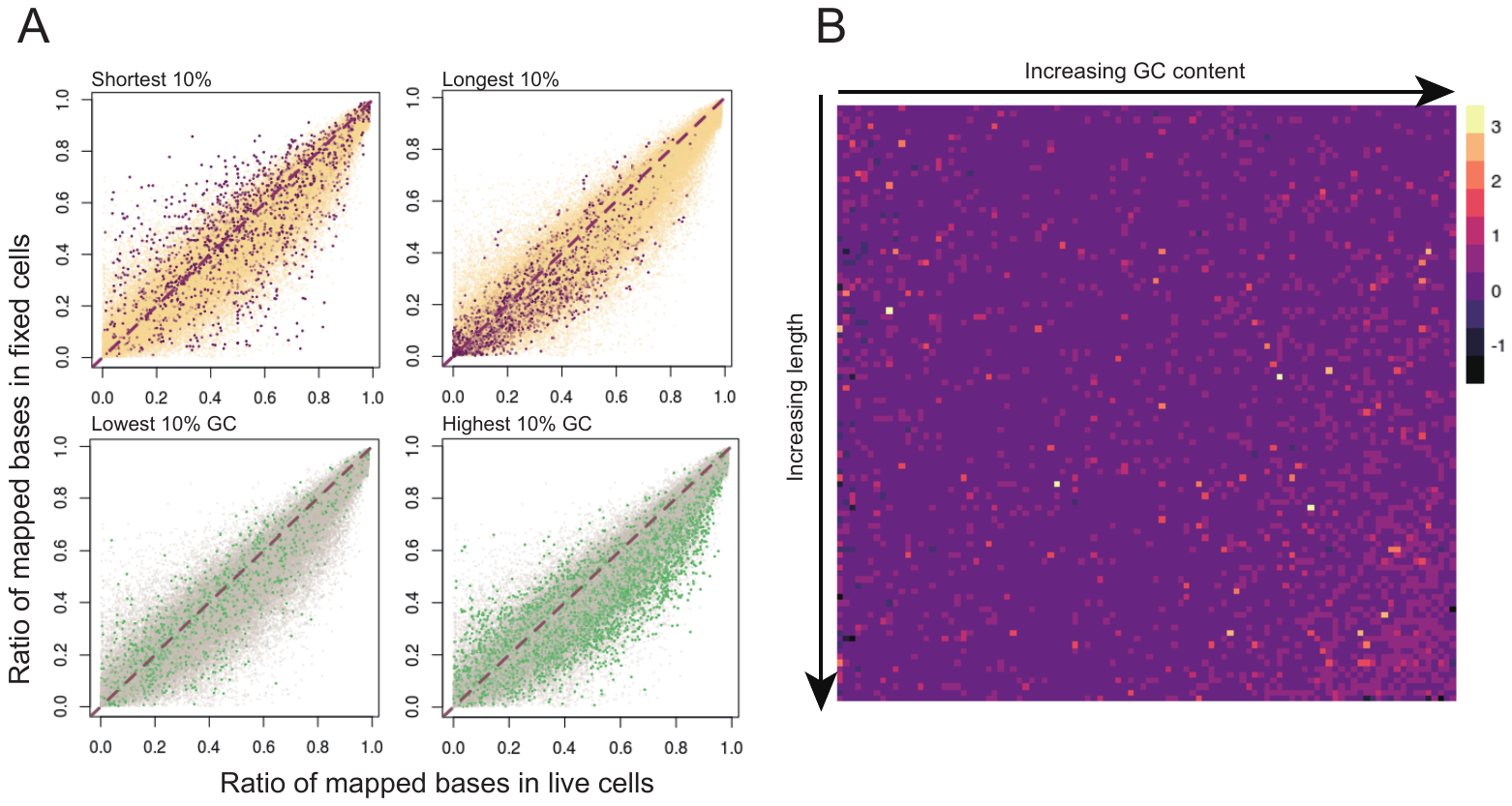

**Supplementary Figure 3.** Comparison of mapping features between live and fixed cells in HepG2.

**(A)** The mapping ratios for each transcript were compared using coverage integrity correlation. Transcripts with top or bottom 10% rank in length and GC content are highlighted in each correlation plot.

**(B)** Visualization of the ratio of live/fixed mapping integrity showing transcripts with better coverage. Transcripts are sorted and grouped by length and GC content; each unit represents an average ratio for those transcripts

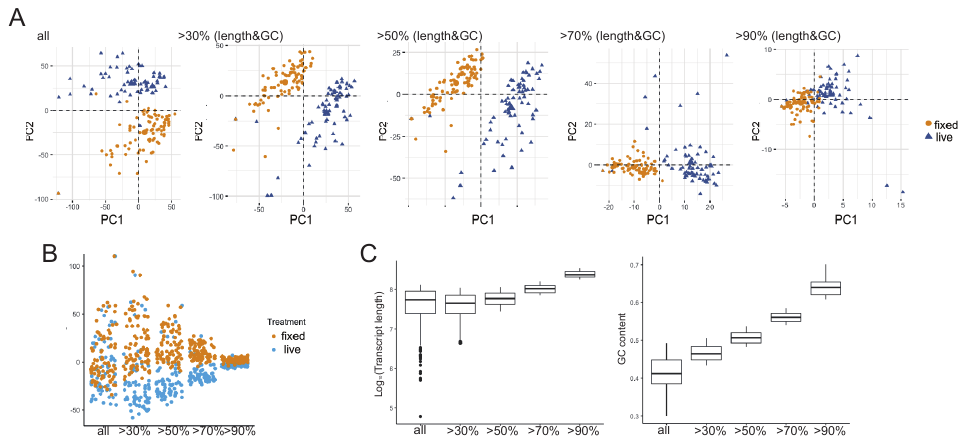

**Supplementary Figure 4.** Transcripts with longer lengths and higher GC contents separate live and fixed cells (done using HepG2).

(**A**) PCA performed using different transcripts sets. In each plot, transcripts are selected based on lengths and GC contents thresholds.

(**B**) PC1 loadings of cells in PCAs performed with different transcripts sets.

(**C**) With PCAs performed with increasing lengths and GC contents thresholds, corresponding length or GC content statistic of the top 500 transcripts from PC1s was plotted.

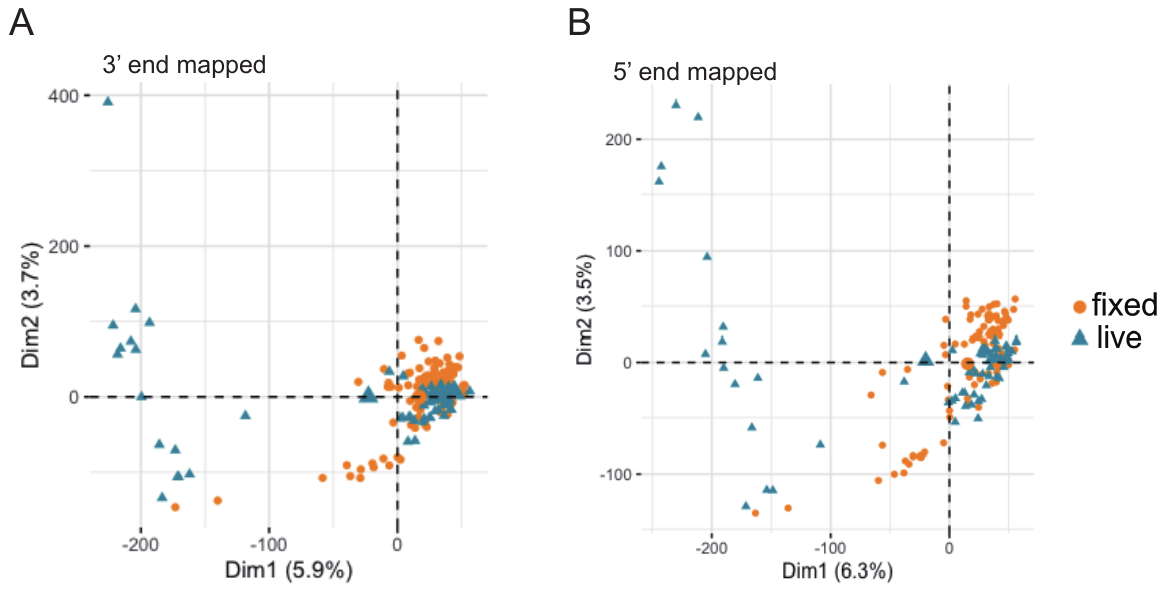

**Supplementary Figure 5.** PCA using HepG2 data generated by mapping raw reads to 3’end (A) and 5’end (B) of transcripts.

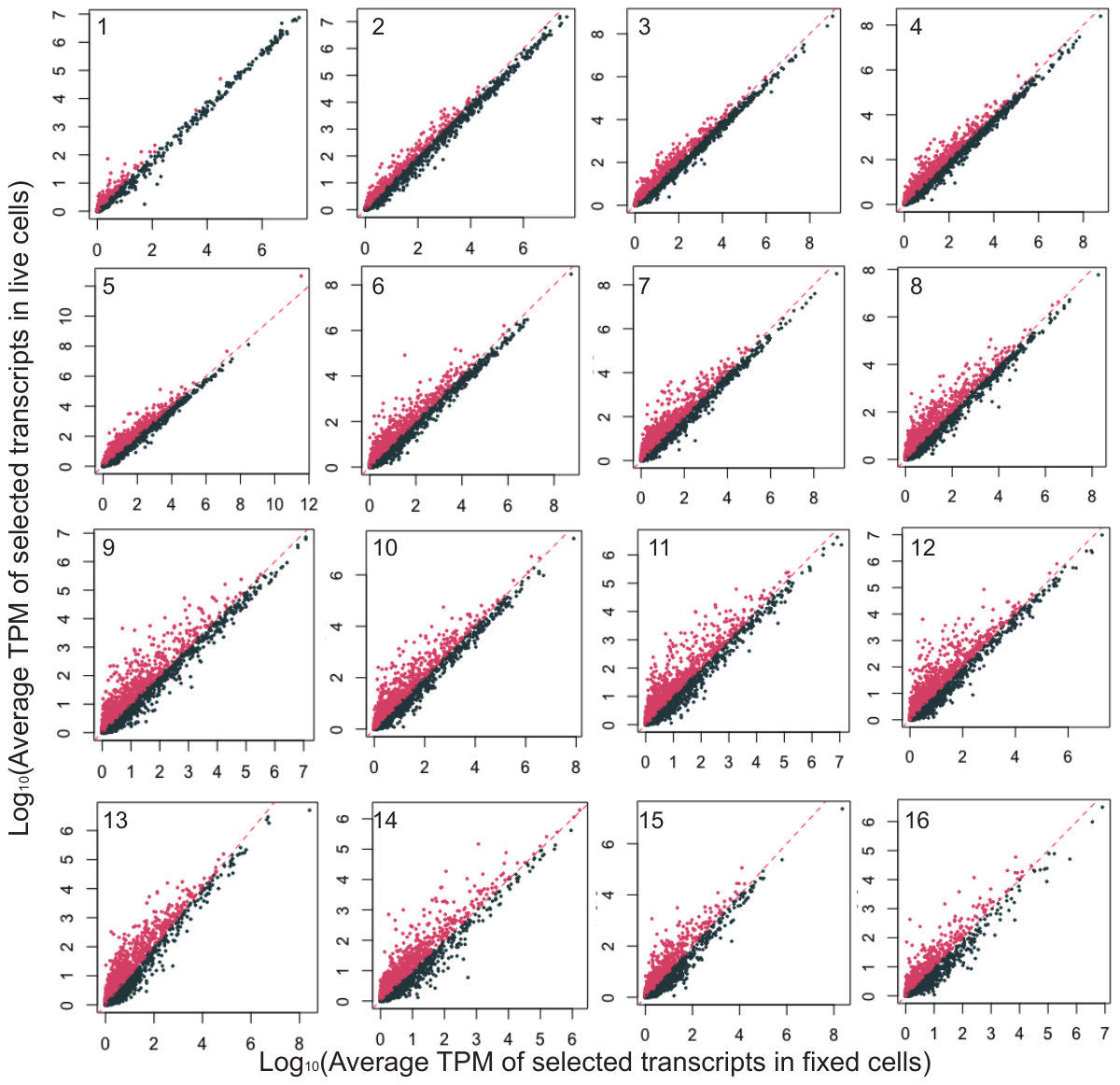
HCT-116

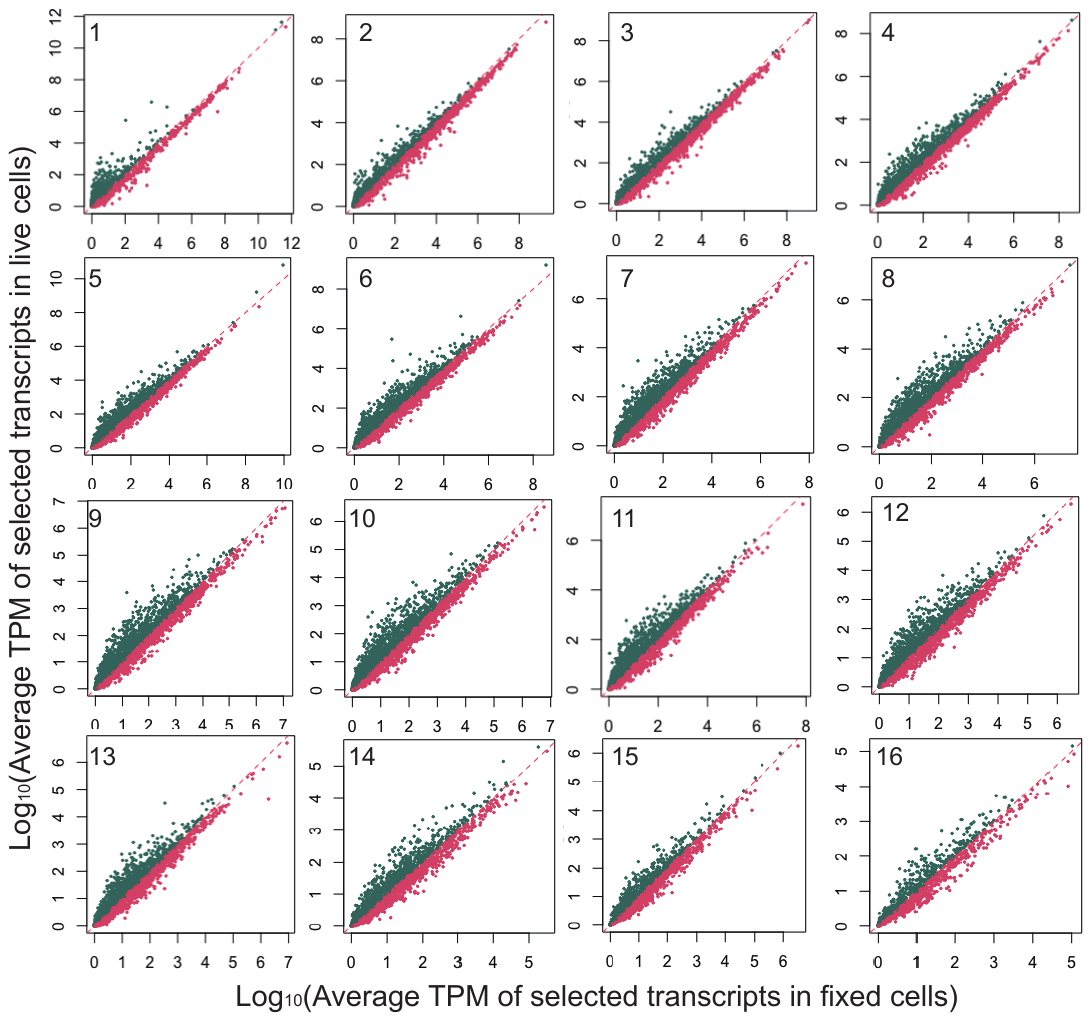

HepG2

**Supplementary Figure 6.** Expression correlation of transcripts with different lengths. Transcripts are equally grouped to 16 according to the length. IDs in each plot represent the transcripts group included in that plot. With the increase of the ID number, the average length of the transcript group also increased. Both results of HCT-116(top) and HepG2(bottom) are shown.

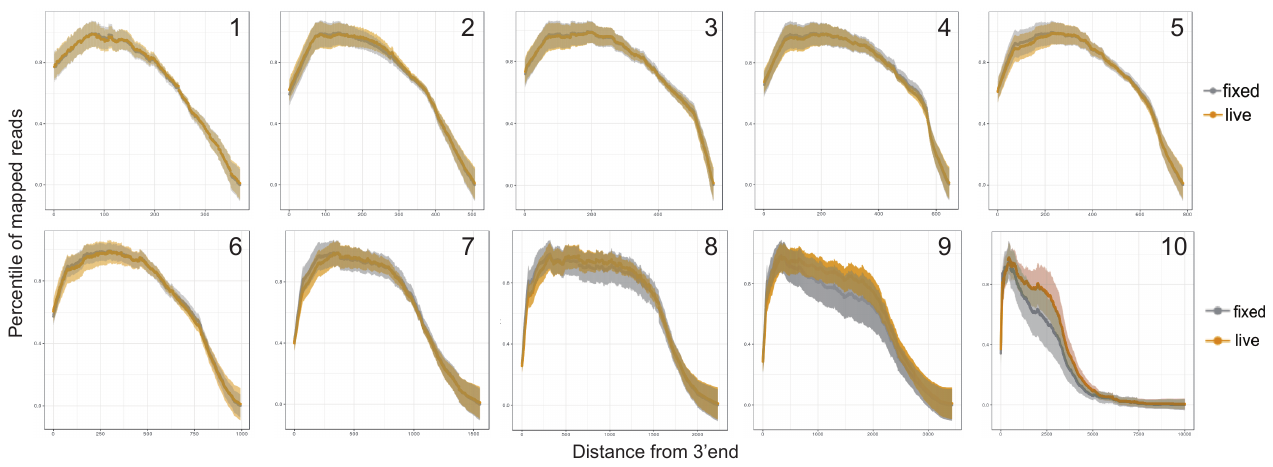
HCT-116

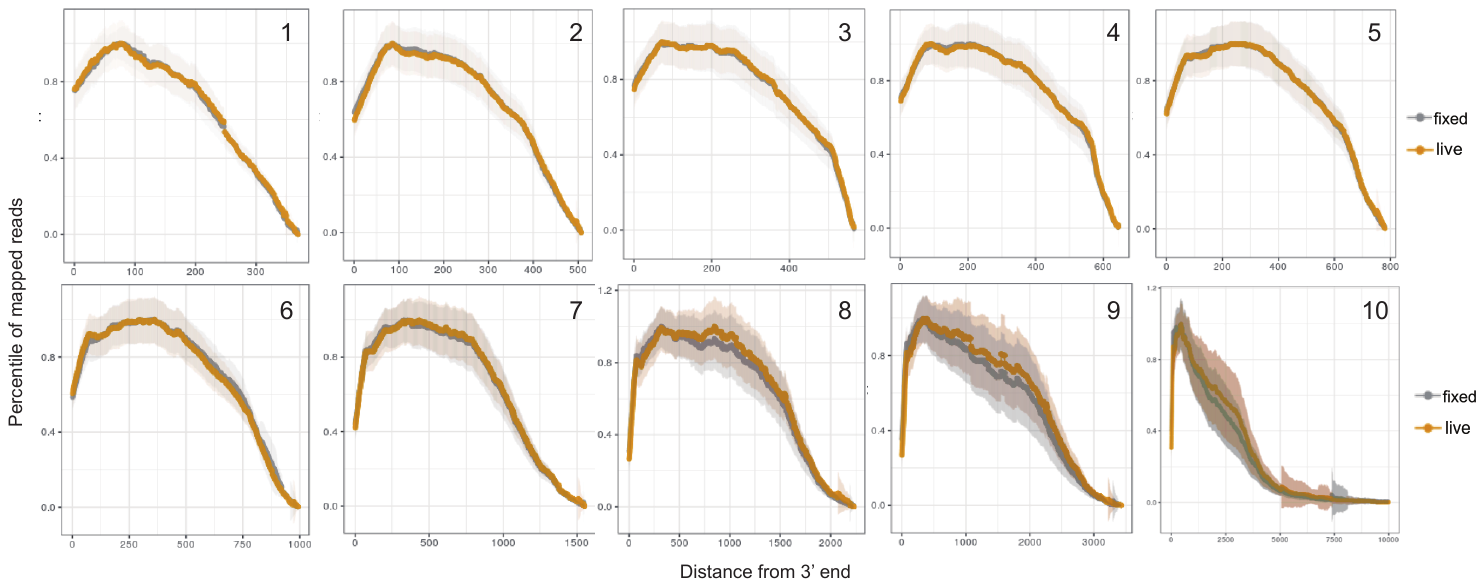

HepG2

**Supplementary Figure 7.** Comparison of mapping coverage between live and fixed cells. Transcripts are equally grouped to 10 according to the length. IDs in each plot represent the transcripts group included in that plot. With the increase of the ID number, the average length of that transcript group also increased. Both results of HCT-116(top) and HepG2(bottom) are shown.
